# Geometrical structure of perceptual color space: mental representations and adaptation invariance

**DOI:** 10.1101/447516

**Authors:** Robert J Ennis, Qasim Zaidi

## Abstract

A central issue in neuroscience is to understand how the brain builds structured representations of percepts that facilitate useful inferences about the world. Similarity between percepts is used to accomplish many everyday tasks, e.g. object identification, so is widely used to construct geometrical spaces that represent stimulus qualities, but the intrinsic validity of the geometry is not tested critically. We introduce an experimental approach to equating relative similarities by setting perceived midpoints between pairs of stimuli. Midpoint settings are used with Varignon’s Theorem to test the intrinsic geometry of a representation space, and its mapping to a physical space of stimuli. For perceptual color space, we demonstrate that geometrical structure depends on the mental representation used in judging similarity: an affine geometry is valid only when observers use an opponent-color mental representation. An affine geometry implies that similarity can be judged within straight lines and across parallel lines, and its neural coding could involve ratios of responses. We show that this perceptual space is invariant to changes in illumination color, providing a formal justification to generalize to all of color space, color constancy results measured for color categories. Our midpoint measurements deviate significantly from midpoints in the “uniform” color spaces CIELAB and CIELUV, used extensively in industry and research, so these spaces do not provide adequate metric representation of perceived colors. Our paradigm can thus test for intrinsic geometrical assumptions underlying the representation space for many perceptual modalities, and for the extrinsic perceptual geometry of the space of physical stimuli.

**Significance:** Mathematical spaces based on similarity judgments are widely used to represent stimulus qualities in perception, cognition and neuroscience. We introduce a perceptual approach to equate relative similarities, and use them to test the geometry of a perceptual space and its mapping to a physical space of stimuli. For color perception, our results show that perceptual geometry depends on the mental representation used in judging similarity, and it has an affine structure when observers use an opponent-color representation. An affine geometry implies that neural coding of similarity could involve simple ratios of responses. Our measurements also reveal that the uniform color spaces CIELAB and CIELUV, used extensively in industrial applications, do not provide adequate representation of similarity between moderately spaced colors.

## Introduction

Sensory organs provide organisms with clues about the environment, but the important properties of the world are rarely explicit in the sensory input. To facilitate dealing with the environment, populations of neurons build and manipulate representations that make explicit useful properties of materials, objects, illuminations, atmospheres, etc. In this paper we examine fundamental properties of these representations, known as perceptual spaces. The methods we devise are of general applicability, but this paper concentrates on color perception.

A perceptual space consists of a set of relevant stimuli along with a set of similarity relationships. Perceptual spaces have been constructed for features as diverse as gloss (1, 2), patterns (3), timbre (4, 5), vowels (6), sound textures (7), gestures (8), biological motion (9), tactile textures (10), tactile orientation (11), odors (12), and others (13). The characteristics of every perceptual space, center on two fundamental properties: dimensionality and intrinsic geometry, which are, in turn, consequences of the space’s metric, i.e., the operation that defines similarity. Historically, similarities have been estimated by errors in matches as estimates of just-discriminable differences, thresholds, and numerical ratings. Based on the experimentally determined properties of the similarity measure, the perceptual space can be assigned a well-defined geometry, thus providing access to a large number of theorems that in turn specify implications of the representational structure. These geometries form a natural hierarchy, with more highly structured geometries placing greater demands on the conditions that the metric must satisfy (14, 15). At the top of the hierarchy is Euclidean geometry and its non-Euclidean relatives elliptical and hyperbolic, which allow representing stimuli as vectors, with well-defined sizes and angles. Affine geometry is one step down the hierarchy: it allows for vector representations on arbitrarily scaled axes, thus defining lines and parallelism, but not angle or size. Further down is projective geometry: collinearity and dimension remains defined, but not parallelism. Lower still,with the fewest geometrical requirements is topological space, where proximity is defined but collinearity and dimension are not. Superimposed on this characterization of the intrinsic geometry of the perceptual space is its extrinsic geometry, which is the mapping of the perceptual space onto a physical space of stimuli, characterizing which can provide additional information about neural transformations. In terms of the dimensionality of the space, higher-dimensional representations enable added flexibility in learning and finer-grained qualitative distinctions, but can impose a higher cost on similarity computations.

Intuition based spaces for color have a long history (16, 17), but the paradigmatic example of the empirical analysis of a perceptual space was provided by Maxwell (18). Maxwell was able to embed color matches into the structure of a linear algebra by showing that color matches satisfy the linearity properties of additivity and scalar multiplication, thus allowing for vector operations to predict the results of combining lights of different colors. Although the physical combinations of visible lights that range from 400-700 nm form a space of infinite dimensionality, Maxwell showed that the space of color-matches was only three-dimensional. His key observation was that color-normal observers could match any light by adjusting the intensities of any three spectrally fixed lights, as long as none of the three primary lights was matchable by a combination of the other two. To probe the physiological basis of color space, Maxwell showed that the matches of congenital color defectives were a reduced subset of color-normal matches, so he could derive the spectral sensitivities of the three types of cones using the additional color confusions of color-blind individuals. Electrophysiological measurements of the spectral sensitivities of human cones (19) confirmed psychophysical estimates derived by Maxwell’s methods (20). Maxwellian spaces based on metamers (physically distinct stimuli that appear identical), e.g. CIEXYZ1931 and Macleod-Boynton (21), have proven invaluable for psychophysical and physiological investigations of the visual system. However, they do not provide a basis for representing similarity because their axes can be scaled arbitrarily without altering metamers, so that relative distances along non-parallel lines are incommensurable.

There have been many attempts to add additional structure to Maxwellian spaces to represent perceived distances between colors. The most systematic attempts have built on MacAdam’s (22) color ellipses. Using an extensive set of data collected on one observer, MacAdam found that errors in making color matches were roughly elliptical in shape and their orientation and size changed systematically in CIEXYZ space. Under the assumption that every ellipse represents a unit of perceived color difference, a transform of CIEXYZ axes that turns all ellipses into circles of similar radius, imposes an isotropic Euclidean metric. Theoretically motivated approximations were proposed by Le Grand (23) and Friele (24), but industry relies heavily on “uniform” color spaces such as CIELAB (25) and CIELUV (25), which use nonlinear transformations based on empirical criterion. All of these spaces suffer from the limitation that each ellipse estimates just noticeable color differences around each local color, without accounting for nonlinear adaptation shifts between separated colors (26). “Uniform” color spaces are often used to choose sets of equally spaced colors spanning color space for psychophyics and electrophysiology experiments, but such sets have not been critically tested against psychophysically measured perceived similarities, as we do below.

The uses of color to make inferences about the environment, and guide action, often require more sophisticated processing than just color-matching and discrimination, e.g. the use of relative color similarities helps to identify materials across spectrally distinct illuminations (27-29). Multidimensional scaling (30) has been used to specify Euclidean color spaces based on numerical ratings of similarity between colors (31), with all the complications of mapping a perceptual quality to numbers. In addition, Wuerger et al (32) showed that proximity judgments between colors fail Euclidean assumptions. Thus, the intrinsic geometry of color space remains to be determined: it should have enough structure to support judgments of relative similarity, but proximity judgments indicate that it is not Euclidean.

We introduce a method to directly investigate the geometrical structure supporting a color similarity space. Varignon’s Theorem (33) states that the bimedians of a quadrilateral bisect each other, i.e. the point of intersection of the two straight lines joining the midpoints of opposite sides is the midpoint of both lines (Figure S1). This theorem holds only for geometrical spaces that are at least Affine. If the vertices of the quadrilateral can be expressed as vectors, then the overlapping mid-points provide two different ways of estimating the same centroid vector (Figure S2). So if Varignon’s theorem does not hold, then the space is not Affine. We tested its validity for perceived centroids of quadrilaterals covering extended areas of color space. Observers viewed a test patch flanked by two patches, each containing one vertex color of the test quadrilateral. They were instructed that a mid-point between two colors is the color that is simultaneously most similar to the two, and could be ascertained by first finding the set of stimuli that are equally similar to two fixed stimuli, and then from this set, the stimulus that is most similar to both. After finding the midpoints for the four sides, observers set the midpoints between the two pairs of facing midpoints. The final mid-point settings were not close to each other for any observer in any condition, thus refuting the Affine assumption. Since color judgments based on “reddish-greenish” and “bluish-yellowish” opponent-dimensions give very stable estimates of color categories (34, 35), observers were then instructed to consider the color difference between the endpoints along the opponent dimensions, and to adjust the middle patch’s hue and saturation to a color perceived as the midpoint on both dimensions, i.e. equally and most similar to both endpoints. For all observers, the two final midpoints for each quadrilateral coincided, thus satisfying the conditions for an Affine space. Therefore, when observers explicitly use an opponent-color mental representation, the perceptual color space of relative similarities across large color differences has an Affine structure. In a Euclidean color space, distance represents magnitude of similarity, but even in a weaker Affine space, ratios of distances along every color line provide measures of relative similarity, and parallelism provides similarity between color changes.

To be widely useful, a neural representation has to be invariant across conditions, just as some extra-striate neurons have object sensitivities that are invariant to pose (36). We show that the geometrical space constructed with the mid-point settings is invariant across different overlaid colored illuminants, and this has significant implications for color constancy (28, 35).

### Experiment 1: Geometrical test of structure for perceptual color space

We used Varignon’s Theorem to test if perceptual color space is Affine by translating it into a series of psychophysical midpoint judgments (See Supplemental Methods). After adapting to a mid-grey background, three rectangles appeared on a calibrated color monitor (Fig 1A). The colors of the two outside rectangles were set to two adjacent vertices of a quadrilateral in the MacLeod-Boynton (21) equiluminant color plane, and observers were instructed to use a joystick to set the color of the central rectangle to the perceptual midpoint of the end-point colors by finding the color that was the most similar to both end-point colors, out of all colors equally similar to both. This was repeated for the four sides defining each color quadrilateral. Then, the mean settings for opposite sides were used as endpoints, and observers were instructed to find the perceptual midpoint. The two midpoints for the two pairs of these end-points are both estimates of the centroid vector, if the four vertices of the quadrilateral can be represented as vectors in an affine space. Therefore, if the final two midpoint settings did not coincide, an intrinsic Affine geometry was rejected for the perceptual color space. Means and standard deviations of midpoint settings were calculated for 10 repeated measurements by each of 4 observers, for a large square and a large diamond that were centered at mid-grey and spanned most of the equiluminant color plane displayable on the monitor (Fig 1B). For all observers, and all quadrilaterals, the two centroid-estimating midpoints did not come close to coinciding (Fig 1C), except for one case. Thus, perceptual color space is not Affine when observers set midpoints based purely on judgments of relative color similarity.

**Figure 1.**
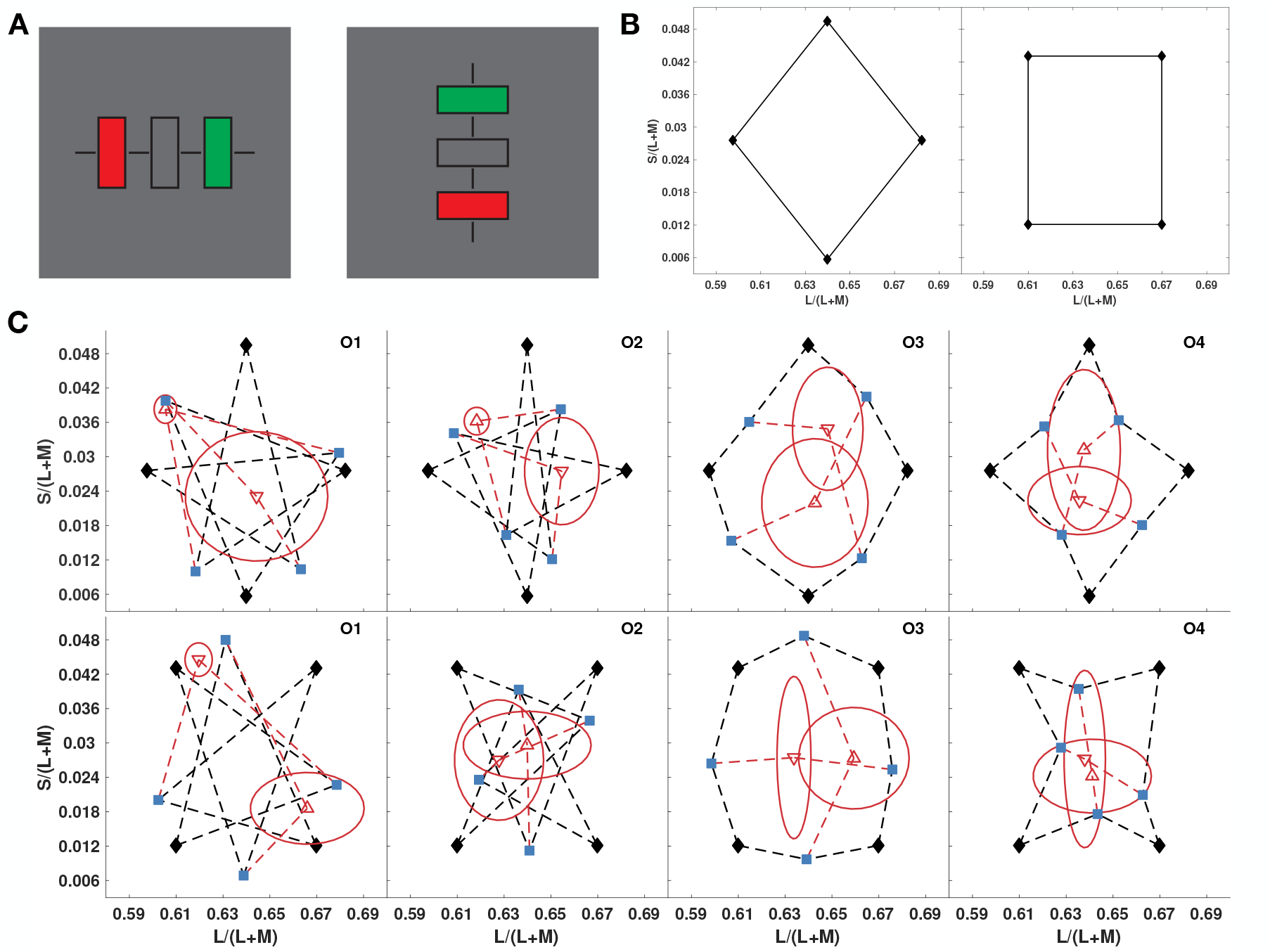
Tests for affine geometry of color similarity (Experiment 1). (A) Stimulus on each trial, alternated between vertical and horizontal on succesive trials, to reduce the effect of aftereffects. Black bars assisted in central fixation. The colors of the flanking rectangles were fixed and observers adjusted the color of the middle rectangle to their estimate of the midpoint color. (B) Color diamond and square used as the quadrilaterals (MacLeod-Boynton chromaticity diagram). (C) Results for 4 observers in Experiment 1 for the diamond (top row, and square (bottom row). Blue points show midpoint settings for the four sides of each quadrilateral, connected to their vertices by black dashed lines. Red points show midpoint settings for opposite blue points, connected by red dashed lines. Red ellipses depict standard deviations for the midpoint settings along the major and minor axes directions.

### Experiment 2: Effect of mental representations on intrinsic geometry

The results of Experiment 1 restrict color similarity space to at best a projective geometry, and this seems to go against observers’ abilities to reliably identify category boundaries between colors based on classifying colors as “reddish” versus “greenish”, and “bluish” versus “yellowish” (35). We thus repeated Experiment 1 with the same observers, but gave them different instructions for finding the perceptual midpoint between the end-point colors: “Consider the colors of the flanking rectangles in terms of ‘Red-Green’ and ‘Blue-Yellow’ qualities. Next, judge the change in each of these qualities between the two colors. Then, set the central rectangle to the color that lies halfway in the ‘Red-Green’ interval defined by the flanking colors, and simultaneously half-way in the ‘Blue-Yellow’ interval.” Observers were given no examples or definitions of “reddish”, “greenish”, “bluish”, or “yellowish”, or of the curvature of the opponent unique hue axes. The results of Experiment 2 in the left two columns of Fig 2 show that the midpoints estimating the centroid were coincident and lie near the center of each quadrilateral. Error ellipses in Experiment 2 for the same quadrilaterals as Experiment 1, are appreciably smaller. Consequently, similarity judgments made with opponent-color mental representations satisfy the conditions for an intrinsic affine geometry.

**Figure 2.**
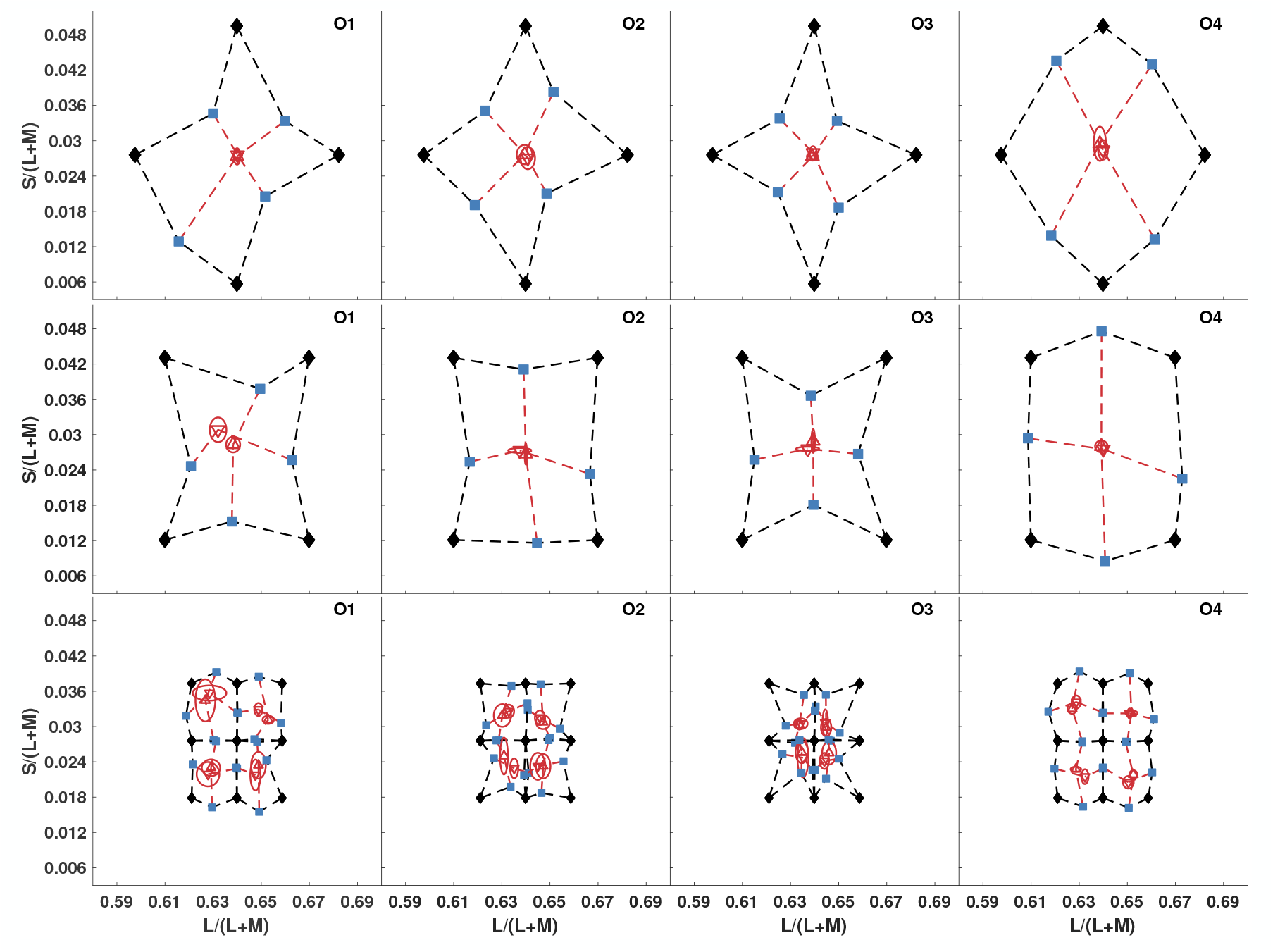
Testing for affine geometry of color similarity using opponent-color mental representations (Experiment 2). We tested the same two color quadrilaterals as in Experiment 1 (top two rows), but with opponent-color instructions. We also tested four additional, smaller quadrilaterals, all of which shared a vertex at mid-gray (bottom row). Same plotting conventions as Fig. 1C.

To further test the affine nature of the space, we also used four smaller squares, each of which had one vertex on mid-gray, instead of being centered on it (Fig. 2 bottom row). The centroid estimating midpoints were again coincident within small errors, indicating that the intrinsic geometry is affine for mid-point judgments on both the large and small quadrilaterals.

### Experiment 3: Invariance of intrinsic color geometry to adaptation

If the geometric structure of color similarities revealed by Experiment 2 is an efficient representation of functionally important properties, we could expect it to be invariant to adaptation under different illuminations. We repeated Experiment 2, but with the stimuli illuminated by an additional light source. We used a Planar consisting of two LCD displays at 110° angle, superimposed via a 50/50 beam-splitter (Fig 3A). Two medium-sized color quadrilaterals were tested in this experiment, with one of 9 adapting illuminants superimposed on the image of the test stimulus (Fig 3B). Fig 3B right shows the coordinates of the test quadrilaterals with the added light from each of the illuminants. Superimposing the illumination from the top monitor on the stimulus from the bottom monitor is equivalent to adding the illuminant color vector to each test color. All of the test vertices and the illuminants had the same luminance, so in Macleod-Boynton chromaticity space, this shifts the quadrilaterals towards the illuminants and distorts them slightly. Since we were mainly interested in the transformations of perceptual color space that occur under different states of adaptation, observers set just the midpoints of the quadrilateral sides, adapting to one illuminant per experimental session.

**Figure 3.**
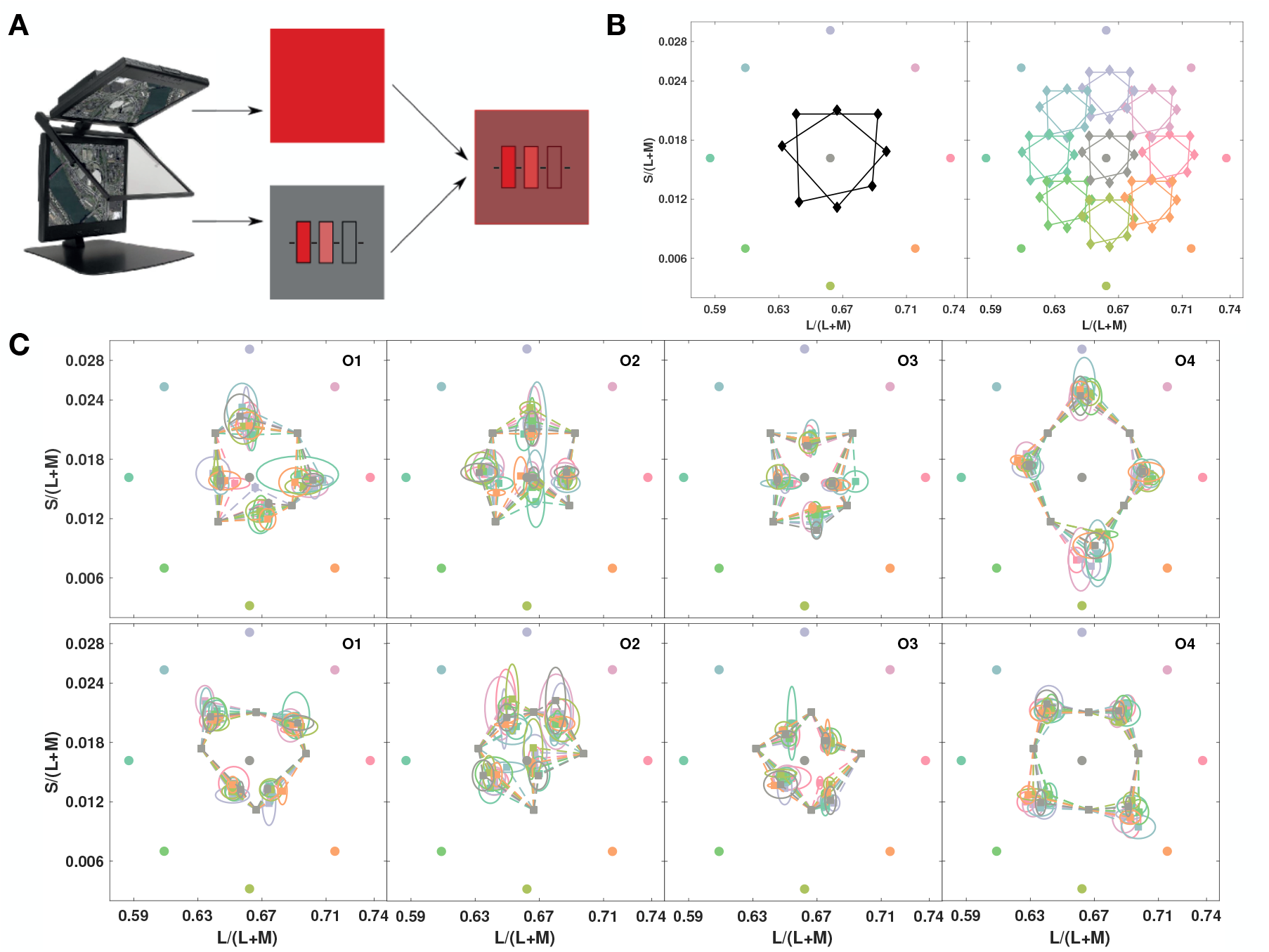
Testing for invariance of relative color similarities across adaptation conditions (Experiment 3). (A) Planar display with beam-splitter superimposing lights from two screens: a full-field red in the top display is combined with the test stimulus in the bottom, allowing test stimulus and task to stay unchanged under different adaptation conditions. (B left) Chromaticities of the nine different superimposed full fields (in roughly corresponding colors) with the tested color quadrilaterals. (B right) Chromaticity coordinates of the lights from test quadrilaterals superimposed with different full-field illuminants (in roughly corresponding colors). (C) Results plotted in the chromaticities of the test monitor for all full-field illuminants. Observers’ chosen midpoints are shown colorcoded according to the illuminant under which they were measured and connected to their respective test vertices, top row for the square, and bottom row for the diamond. Ellipses show the standard deviation of settings, color-coded for the full-field light.

Fig 3C shows the means of 5 repeated midpoint settings for each of nine different states of adaptation. The color of each midpoint symbol indicates the adapting illuminant used for that setting. For all observers, the different adapting illuminants had no appreciable influence on midpoint settings for either of the two tested quadrilaterals. Since the midpoint settings show little variation with adaptation state, their invariance can be explained by a subtractive adaptation process (see Discussion).

## Discussion

We demonstrate that the intrinsic geometrical structure of perceptual color space depends on the mental representation employed by an observer. Goodman (39) pointed out that there are innumerable ways to assess similarity between two real or abstract entities, so the estimated degree of similarity depends entirely on the observer’s perspective. Our results show that this observation also applies to color similarity. If observers judge similarity without being given an explicit representation, then colors seem to aggregate in categories without a natural ordering. This is not entirely unsurprising, because for somebody not schooled in color structure, there is no reason why white and green would not be judged as equally dissimilar colors from red. Instructions to use a color-opponent based representation revealed a perceptual color space with an Affine structure. This allows us to compute ratios between segments of lines to estimate relative similarity between three colors on a line, and to predict observers’ responses to parallel color changes, as would happen to colors of objects across different colored illuminants (40). The main effect of an opponent mental representation seems to be to locate white or grey in the center of the perceptual space, roughly at the intersection of the opponent dimensions. It is worth reiterating that “reddish”, “greenish”, “bluish” and “yellowish” were defined by each observer only mentally and individually. The role of unique hues has been debated because there is no psychophysics or physiology showing primacy for the four unique hues, red, green, blue and yellow (41). The results of this study suggest that they provide a possible systematic arrangement of colors, that our observers were able to use almost effortlessly, much like a mental representation of the cardinal compass directions. Whether this would also be true for speakers of languages that do not have these color terms remains to be seen.

This study was not designed to address the extrinsic geometry (curvature) of perceptual color space, but the midpoints also provide pairs of equal color similarities, so the chromaticities of the midpoints provide clues to the sign of curvature. A quadrilateral on a flat surface, would bulge out if mapped on an ellipsoid, but on a hyperbolic surface, the sides of the quadrilateral would bend inwards and the internal angles would be less than 90°. With the opponent mental representation, the perceived midpoints of the quadrilateral sides lie consistently closer to the center for three observers, indicating an extrinsic hyperbolic geometry. One observer’s mid-point settings, however, bulge outwards. Generally, each observer’s midpoints for each side of the quadrilateral follow a similar pattern as for the large quadrilaterals, suggesting the same class of extrinsic geometries for similarities across large versus small color separations. It seems that extrinsic geometries could be different across observers even if intrinsic geometry is the same. It may thus not be possible for a single nonlinear transform of a chromaticity space to represent similarities for all observers.

To test how well commonly used “uniform” color spaces, CIELUV and CIELAB, represent color similarities across the distances we tested, we plotted observers’ mid-point settings in these spaces (Fig 4). In a truly uniform space, the perceived and calculated mid-points should coincide. For the large square and diamond quadrilaterals in Experiment 2, T-tests (p < 0.05) for 4 observers and 8 midpoint settings revealed that 18 out of 32 measured midpoints deviated significantly from predicted midpoints in CIELAB, as did 18 out of 32 in CIELUV. Across the smaller quadrilaterals, the empirical midpoints were significantly different from predicted for 47 out of 128 in CIELAB, and 33 out of 128 in CIELUV. Average departures of the midpoints for the larger quadrilaterals from the predictions were 5.2±2.49 (mean±SD) in DE units in CIELAB according to CIEDE2000, and 15.67±6.67 (mean±SD) in DE units in CIELUV. Thus, neither of these “uniform” spaces does an adequate job of representing color similarities between separated colors, and thus neither transform from CIE space provides an adequate representation of the extrinsic geometry of the color space. These spaces are probably adequate for delimiting industrial tolerance of color specifications, but should not be used to estimate perceptual distances of separated colors, or to define a set of separated equally spaced colors.

**Figure 4.**
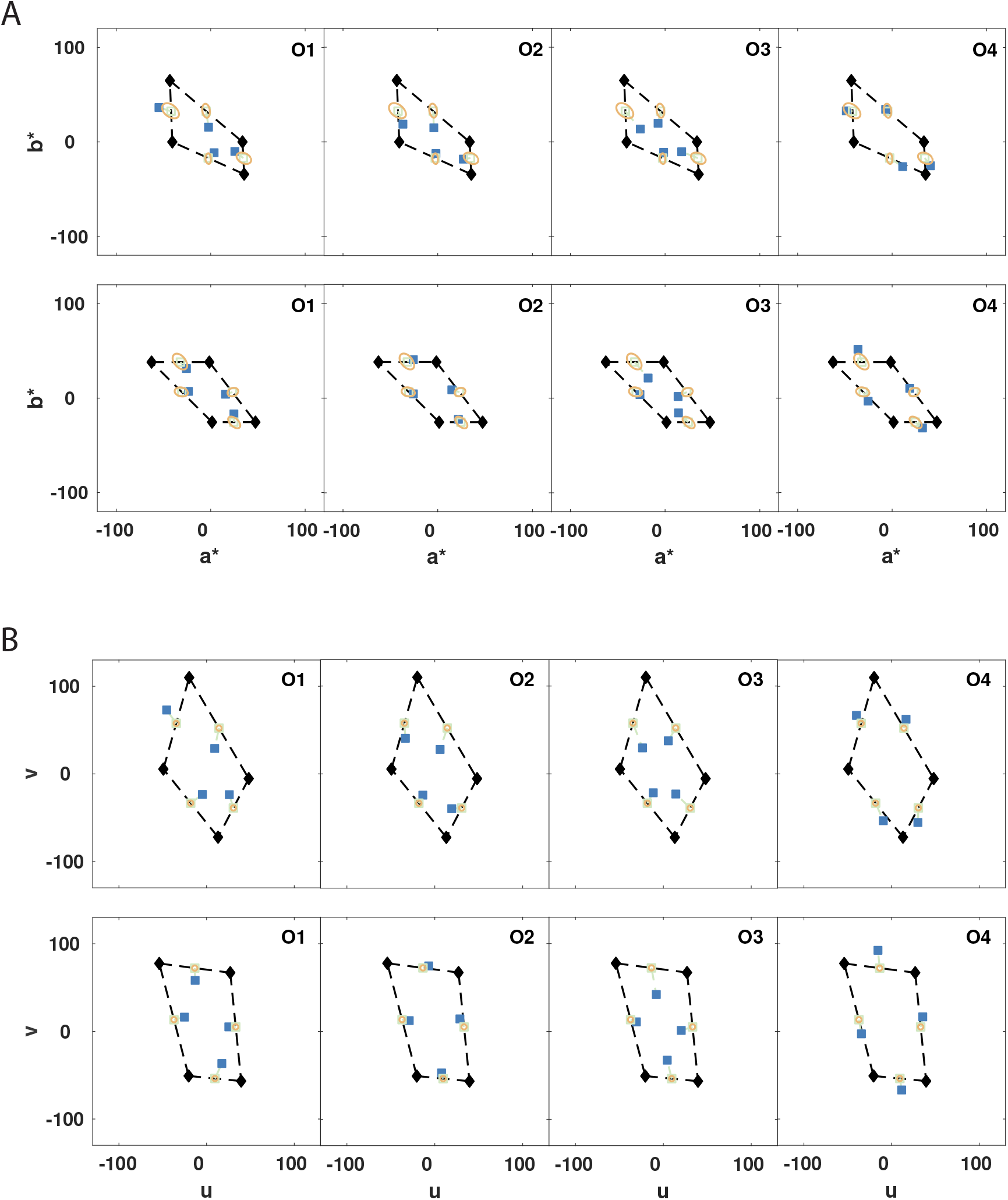
“Uniform” color spaces tested for perceived mid-points, CIELAB (A) and CIELUV (B). The data are from Fig. 1B, square top rows, and the diamond in bottom rows. The shapes are distorted by the non-linear transformations of CIELAB and CIELUV. Blue squares show the average midpoint settings from each observer for the respective edge. The green empty squares show the predicted midpoint between the vertices. Thin green dashed lines connect the predicted midpoint with the blue setting of each observer. The yellow ellipses are 3 DE units, calculated according to CIEDE2000 in CIELAB and the standard Euclidean distance in CIELUV, and are centered on the predicted, green midpoints. Results in CIELAB (a* vs b*), and CIELUV (u vs v), and the statistical analyses in the main text show midpoint settings do not coincide with predicted midpoints.

Adapting to added full-field illuminants distributed about the hue circle did not affect midpoint settings, as if the visual system completely discounted the illuminant by subtracting it from the mixed stimulus. Superimposing the illuminant just creates an additive shift in the colors of the end-points, and this shift retains the shape of each quadrilateral. Hence the discounting is reminiscent of Bergström’s (42) illumination and color analysis, which proposed that the visual system finds the common color vector across surfaces, and segments it from the scene to find the relative components of each reflecting surface. Although the process is compatible with a central process, it is more likely that the two minutes of adaptation prior to the measurements were sufficient to evoke automatic subtractive adaptation mechanisms in ganglion cells of the retina (43), which counteract the additive shift.

The adaptation invariance of geometrical color space provides a formal justification to generalize the color constancy measurements of Smithson and Zaidi (35) to all of color space. They studied color constancy of patches simulating object colors in a variegated background under simulated Sunlight and Skylight. Completely adapted observers were asked to state whether the patch appeared “reddish” or “greenish”, and “bluish” or “yellowish”, thus providing estimates of color category boundaries. Plotted in terms of object reflectance, the category boundaries were found to be invariant to illumination change, indicating a high level of color constancy when observers are completely adapted to a single illuminant. However, invariance of object colors on category boundaries does not guarantee invariance of object colors within boundaries. The invariance of midpoint settings under different adaptation states in this study, establishes that if locations of points on category boundaries are invariant, then since every interior point could be expressed as the midpoint of two boundary points, its location would also be invariant.

Observer mid-point settings were first used by Plateau (37) to informally show that the grey mid-point between white and black was essentially invariant to illumination and observer. This result restricts any hypothetical psychophysical scale that takes intensities to real numbers, to be either a power or logarithmic function (38). A mid-point setting equates two similarities, and we show that color mid-point settings provide consistent and reliable estimates of relative similarity, and these are also invariant to illumination color. However, perceptual dimensions of color, such as hue, saturation and brightness, are not independent, so deriving psychophysical scales for these dimensions requires considering their interactions.

At an abstract level, similarity is one of the fundamental principles used by Gestalt psychology to explain perceptual organization (44), and can be used to generate models of generalization ranging from set-theoretic (45) to continuous metric-space structures (46), especially within a Bayesian formulation (47). Mid-point settings, representing equal relative similarities, can test the intrinsic geometry of a perceptual space as a logical preliminary to using multidimensional scaling, as they test the validity of the assumptions inherent in the statistical procedure. The methods in this paper are easily applied to other modalities, and could thus be used to critically test the geometrical structure of perceptual spaces that have been proposed on the basis of multi-dimensional scaling analyses for many other attributes, such as gloss, timbre, vowels, gestures, biological motion, tactile textures, tactile orientation, odors, and others (13).

Despite the extensive theoretical and empirical work on perceptual similarity, the neural basis of similarity computations is essentially unknown. Possibilities include activation patterns of receptors or later neurons, in rates or temporal patterns of impulse responses, and in different levels of correlated firing. The Affine geometrical structure we identify for color similarities suggests a simplification for neural circuits that compute similarity. Whereas calculating Euclidean distances in a perceptual space implies comparisons based on the power (sum of squares) of the difference, Affine geometry implies simpler comparisons based on ratios. It is possible that perceived colors are decoded using winner-take-all schemes on the responses of individual color-tuned IT neurons (48). In that case, analyses of the population responses of IT cells may help us understand whether the restriction of perceptual color space to an affine geometry, represents a trade-off between the goals of providing an efficient representation of sensory stimuli, and the costs of neural computations.

## Methods (Supplement contains more details)

### Observers

_Four color normal male observers, aged 27-32, gave written consent. The research was approved by the SUNY Optometry IRB.

### Data Analyses

Means and standard deviations of midpoint settings were the key statistics. Ellipses shown on the figures represent +/-1 SD.

#### Experiments 1 & 2

Stimuli were presented on a calibrated HP1230 CRT (Hewlett-Packard, Inc.; Palo Alto, CA, USA), driven at 85Hz by a Visage (CRS, Ltd.; Kent, UK) at 12 bits/gun. Colors in the MacLeod-Boynton (21) chromaticity diagram were displayed using the procedure in Zaidi and Halevy (49). Observers used an Xbox 360 controller (Microsoft, Corp.; Redmond, WA, USA) to set the target color. Three parallel 4° x 0. 6° rectangles (Fig. 1A) were placed at 0.6° separations on a mid-gray background. Each experiment was split into blocks of 20 trials each. There were 2 minutes of adaptation to the mean gray background before the first trial, and 2 second re-adaptation after each trial. The pairs of flanking colors on each trial were chosen by randomly sampling adjacent vertices from the tested quadrilaterals.

#### Experiment 3

Stimuli were presented on a Planar SD2620W (Planar Systems, Inc.; Hillsboro, OR, USA) with two LCD displays at an angle of 110° superimposed by a bean-splitter (Fig 3A), each driven independently at 60Hz by a dedicated port of an NVidia Geforce GTX 580 (NVidia, Corp.; Santa Clara, CA, USA). The displays had already been corrected for spatial distortions, color purity, and alignment (50) and were calibrated through the beam-splitter. The top monitor was used as a full-field spatially uniform illuminant and its image was superimposed on the rectangles from the bottom monitor, simulating a lit scene. One of 9 adapting illuminants was randomly chosen for each experimental session.

## Supplemental information

**Figure S1.**
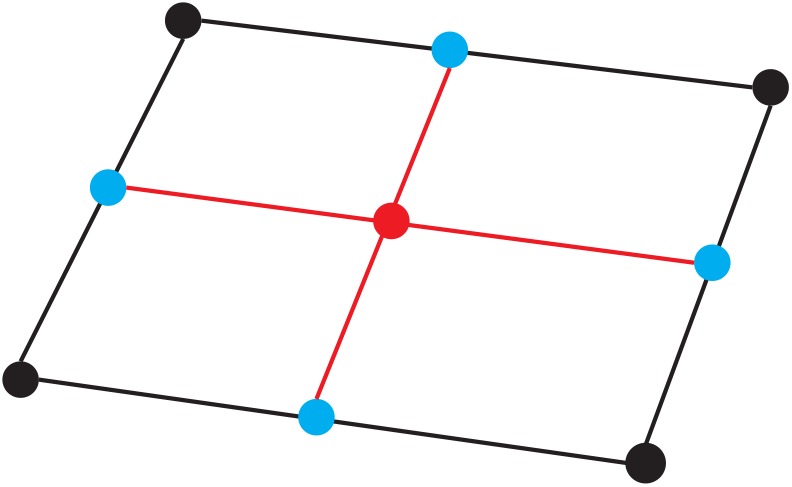
Varignon’s Theorem and Affine Space. Varignon’s Theorem illustrated for an arbitrary quadrilateral. Varignon’s Theorem states that for any quadrilateral, the figure formed by the mid-points of the four sides is a parallelogram. Since the diagonals of a parallelogram intersect at their midpoints, and the diagonals of the parallelogram are the bimedians of the quadrilateral, the point of intersection of the bimedians is simultaneously the midpoint of both lines. The chance of two random points coinciding is exceedingly small, hence the importance of this discovery. However, the theorem became easy to prove after the invention of vector algebra. If the vertices of the quadrilateral can be represented as vectors in an affine (or Euclidean) space, the two overlapping mid-points are represented by the same centroid vector, just calculated in different steps. The theorem can thus serve as a test of whether points in the space can be represented as vectors, subject to the rules of vector manipulation, with the non-coincidence of the midpoints refuting the conjecture that the space is at least affine. The black lines and vertices denote the original quadrilateral. The blue points denote the midpoints on the four sides of the quadrilateral and the red points (overlapping) denote the midpoints of the bimedians (red lines).

**Figure S2.**
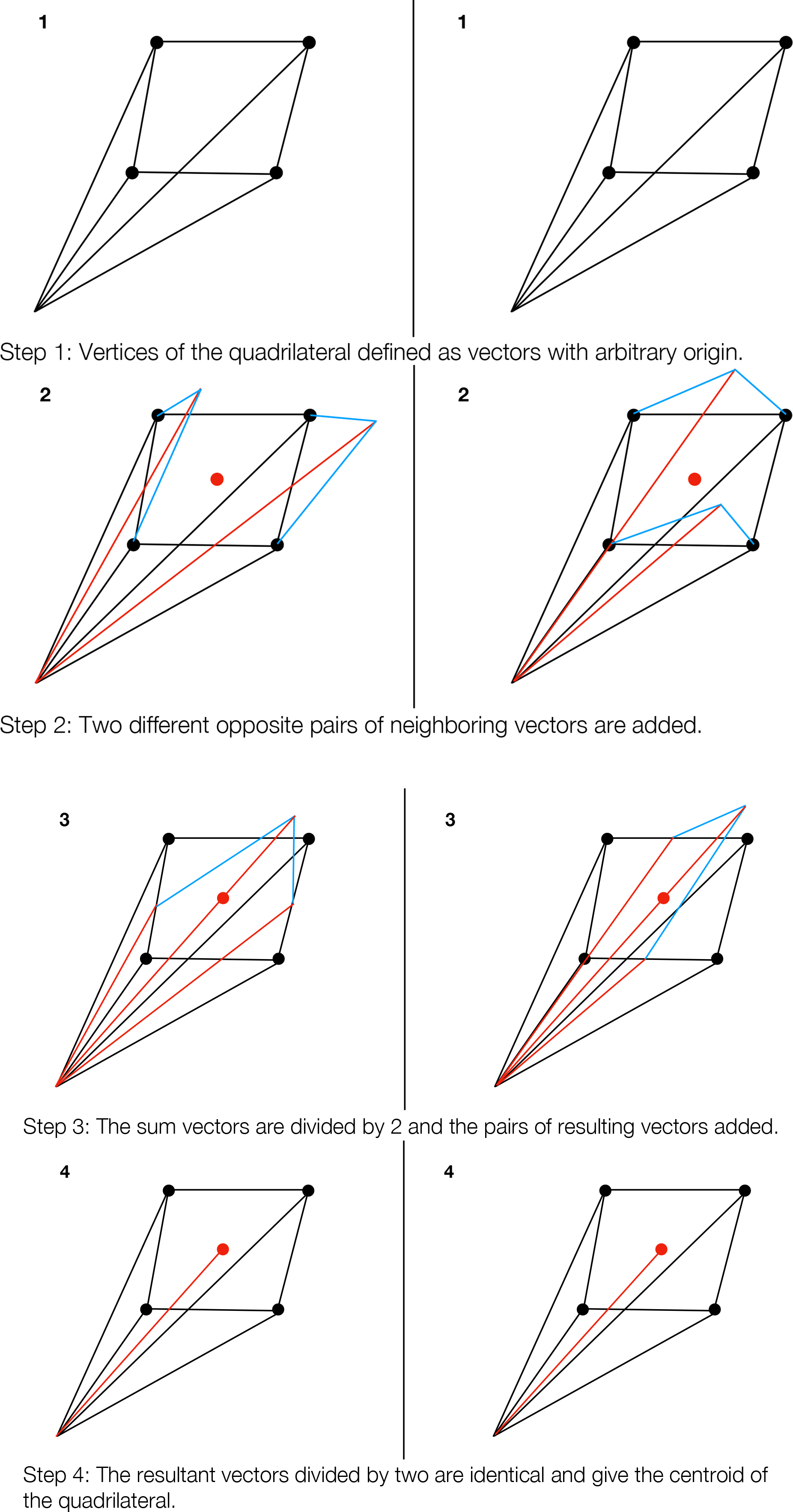
Varignon’s Theorem proven through vector addition: If the vertices of a quadrilateral can be represented as vectors, the midpoints of both bimedians are the same centroid. Consequently, if the mid-points do not coincide, we can conclude that the vertices cannot be represented as vectors in an affine or Euclidean space.

## METHODS

### Observers

Four male observers, aged 27-32, participated in all three experiments. One of the observers was the author, RE. All observers had normal color vision in both eyes. If necessary, observers wore their corrective lenses. All participants were fully briefed on the purpose of the study at the completion of all experiments. The research was approved by and followed the policies of the SUNY Optometry IRB.

### Experiments 1 & 2

#### Equipment

Stimuli were presented on an HP1230 CRT (Hewlett-Packard, Inc.; Palo Alto, CA, USA). The HP1230 was driven at 85Hz by a Visage (CRS, Ltd.; Kent, UK) connected to a DELL Precision 380 (DELL, Inc.; Round Rock, TX, USA) running Windows XP Service Pack 3 (Microsoft, Corp.; Redmond, WA, USA). It had a spatial resolution of 1024×768, minimum luminance of 0.1 cd/m^2 and a maximum luminance of 124.01 cd/m^2. Stimuli for the HP1230 were programmed in MATLAB 2010a (Mathworks, Inc.; Natick, MA, USA) using the CRS Toolbox for MATLAB (v1.27).

Before calibration, we used the monitor’s software menu to tune its settings for an optimal and accurate display of images (convergence, zero curvature of straight lines, color purity, etc.). The luminance curves for each phosphor of the HP1230 was measured using an OptiCAL (CRS, Ltd.; Kent, UK) via the VSG Desktop (v8.200) interface provided by CRS, Ltd. The monitor’s voltage-vs-luminance curves, sampled at 256 levels for each gun, were fit with power curves using a least-squares procedure that was automated by the VSG Desktop. The chromaticity coordinates of the phosphors were measured with each at their maximum using a PR650 spectroradiometer (Photo Research, Inc.; Syracuse, New York, USA). For all stimuli, colors were computed in the MacLeod-Boynton (MB) color space (1, 2) and converted to RGB coordinates via a transformation matrix using the procedure reported in Zaidi & Halvey (3). The CRS Toolbox’s internal routines used the lookup table generated by the VSG Desktop calibration routine to properly transform the input and present the desired MB color on the HP1230.

Observers used an Xbox 360 (Microsoft, Corp.; Redmond, WA, USA) controller to manipulate the target patch in each trial and input their final response. The MATLAB joymex2 package was used to interface with the Xbox controller. The controller had two joysticks. The left joystick was pushed left<->right to change the purity of the target patch and the right joystick was pushed up<->down to change its hue. These corresponded to changing the length and angular position, respectively, of a vector originating at mid-gray in the equiluminant plane of the MB space. When observers were satisfied with their midpoint setting, they pressed the “Y” button on the Xbox controller to continue to the next trial.

#### Stimuli and task

All stimuli were viewed binocularly. The background was set to mid-gray. At the center of the monitor, three parallel 4° x 0.6° rectangles (Fig. 1A) were placed side-by-side at 0.6° separations. The rectangles were either all oriented horizontally or all oriented vertically. The orientation switched on each trial and was not randomized in order to keep a balanced adaptation state across the visual field. The colors of the outermost flanking rectangles were fixed on each trial and were determined by randomly sampling neighboring vertices from color quadrilaterals specified in the equiluminant plane of the MB space (see Figs. 1 and 2 in main text). The color of the central rectangle could be controlled by the observer. The central rectangle was always presented at the center of the screen. Each rectangle had a 0.02° (2px) black border. Four black bars (0.02° x 0.6°) were presented orthogonal to the three rectangles and were vertically aligned along their centers to guide fixation. Otherwise, observers were allowed to freely fixate.

The goal of observers was to change the color of the central target rectangle until it appeared to be the midpoint color of the two flanking colored rectangles. They did this according to different instructions in Experiments 1 and 2. In Experiment 1, observers were instructed to use the joysticks to set the color of the central rectangle to the perceptual midpoint of the end-point colors by finding the color that was the most similar to both end-point colors, out of all colors equally similar to the end-point colors. In Experiment 2, observers were instructed to “consider the colors of the flanking rectangles in terms of ‘Red-Green’ and ‘Blue-Yellow’ qualities. Next, judge the change in each of these qualities between the two colors. Then, set the central rectangle to the color that lies halfway in the ‘Red-Green’ interval defined by the flanking colors and simultaneously half-way in the ‘Blue-Yellow’ interval”.

#### Testing sequence

Each experiment was split into blocks of 20 trials each. The number of blocks per experiment depended on the number of color quadrilaterals, and the number of midpoint measurements. We measured 10 midpoint settings per edge for each quadrilateral and another ten midpoint settings for the bimedians. The first block was preceded by 2 minutes of adaptation to the mean gray background. After every trial, the relevant data were saved and a 2 second pause for re-adaptation to the background preceded the next trial. After every block, the observer was given the choice of either continuing the experiment or stopping. If the observer chose to continue, a one minute re-adaptation period was initiated before the next block of trials began. If the observer chose to stop, the program saved the current state of the experiment. The next time the experiment was run, the program picked up from where the observer had stopped and initiated the next block of trials with 2 minutes of adaptation. The pairs of test colors on each trial were chosen by randomly sampling edges from the tested quadrilaterals. Once an edge was chosen, the two vertices that defined it were used as the colors for the flanking rectangles.

#### Data Analysis

Means and standard deviations of midpoint settings were used as the key statistics. Ellipses shown on the figures in the main text represent the standard deviation of settings along the axes of the MacLeod-Boynton diagram (i.e., the axes of the ellipses are aligned with axes of the diagram and their radius is the standard deviation of the settings in that direction).

### Experiment 3

#### Equipment

Stimuli for experiment 3 were presented on a Planar SD2620W 3D LCD Display (Planar Systems, Inc.; Hillsboro, OR, USA). The Planar had two LCD displays, each of which was independently driven at 60Hz by a dedicated port of an NVidia Geforce GTX 580 (NVidia, Corp.; Santa Clara, CA, USA). The bottom of the two Planar monitors was situated perpendicular to the observer’s line of sight. The top monitor was oriented 110° away from the bottom monitor, and the images that each monitor generated were superimposed via a beamsplitter with a partly reflective coating that bisected the angle formed by the two monitors (fine adjustments were made to eliminate shadows at the edges of the display). Stimuli for the Planar display were programmed in MATLAB 2012a 32-bit (Mathworks, Inc.; Natick, MA, USA) using Psychtoolbox (v3.0.9). The computer was custom built by iBUYPOWER (iBUYPOWER, Inc.; Los Angeles, CA, USA) and ran Windows XP Service Pack 2 64-bit edition (Microsoft, Corp.; Redmond, WA, USA). The image from each of the Planar monitors was polarized for the presentation of 3D stimuli (the polarized light from the top monitor was rotated 90° relative to the polarized light of the bottom monitor), but we did not make use of this feature.

Observers viewed the Planar display without the 3D polarized glasses, since no 3D stimuli were displayed and the glasses would have reduced the overall luminance reaching each eye. Rather, the top monitor of the Planar setup was used as a full-field illuminant and its image was superimposed on the image from the bottom monitor, simulating a lit scene. The view of the top monitor was blocked by a black piece of cardboard to prevent observers from viewing the illuminant and gaining any information about it. A black piece of cardboard was also attached to the bottom of the mirror, since it was possible to view part of the background on the bottom monitor without it. With both pieces of cardboard in place, the view of the stimulus was restricted to the superimposed illuminant-stimulus image.

The outputs of each Planar display were measured after each was filtered through the mirror. The Planar monitors were calibrated with the same PR650 that was used to calibrate the HP1230. Each of the Planar’s two displays were calibrated while the other was turned off. A 1sec delay preceded each measurement with the PR650 to allow the LCD to settle into a stable state. We presented full screen patches on each display and calibrated both to have a similar mid-gray white point (CIE x:0.329, y:0.338) and min and max luminance (min = 0.11 cd/m^2, max = 116 cd/m^2). We tested that the spectra from the primaries for each of the Planar displays overlapped and were indistinguishable by eye. The chromaticity coordinates of the three primaries were recorded at their maximum output and were checked for spatial/temporal stability and equality across the two displays. The chromaticity coordinates of the three guns on the LCD were measured with the PR650 and used to construct a transformation matrix that took coordinates in the MB space to 8-bit RGB values, similar to the transformation matrix for the CRT setup. The Planar displays had already been corrected for spatial distortions, color purity, and alignment in a previous study (4). No deviations from these settings were found during calibration, so the monitors were left untouched. In particular, we ensured that the monitors were set to their factory standard display settings and were not emulating a showroom environment or a movie theater.

The input-output curves of each of the primaries were measured for a square, 6° patch on a mean gray background that was stepped through the entire range of each primary, one at a time, while the other two primaries were set to their mid-level output. This was done to calibrate the LCDs to their outputs about the mid-gray, since LCDs are known to have poor gray tracking (5) and all of our equiluminant stimuli were essentially small-to-medium deviations from the mid-level output of each primary. Luminance measurements were taken with the same OptiCAL used to measure the HP1230, via a MATLAB<->PyOptical interface (6; free, open-source software developed by Valentin Haenel and modified for our lab setup by Martin Giesel). Lookup-tables (LUTs) were generated for each primary input-output curve (8-bit input = 0-255). The input (bit level) vs. output (luminance) relationship for each primary, as measured with the OptiCAL, was fit with a spline via MATLAB’s internal fitting routines. When a specific luminance for each gun was needed, the closest value in the LUT was found and the corresponding bit level was sent to the graphics card for display. The linearity of the gun outputs was verified, after the inputs had been calibrated via the LUT. The LCD input-output curves, after linearization, were fit with lines (all fits had R>0.99). The beam-splitter mirror was tested for linear additivity. We measured the output of each monitor individually at 6 different gray levels (0%, 10%, 25%, 50%, 75%, and 100%) through the mirror. Then, we measured the combination of both monitor outputs at each of these levels through the mirror. The superimposed gray levels showed no deviation from addition of the individually measured outputs. Observers used the two joysticks of the Xbox 360 controller to adjust the hue/saturation of the center rectangle, just as in Experiments 1 and 2.

#### Stimuli and task

For this experiment, observers used the same instructions from Experiment 2 and saw the same stimulus, but now one of 9 adapting illuminants was superimposed on the image of the three rectangles. Since we were solely interested in the transformations of perceptual color space that occur under different states of adaptation, only the midpoints of the sides were measured. The two quadrilaterals in Fig. 3B of the main text were tested with these conditions. Since all observers had enough practice with midpoint settings and midpoint settings were found to have small variability across trials in Experiment 2, only five measurements of each midpoint per edge of a color quadrilateral were taken, giving a total of 180 trials per observer.

#### Testing sequence

For Experiment 3, the adaptation states were randomly sampled, but only one adaptation state was tested during each experimental session. The test pairs for Experiment 3 were then randomly sampled in the same manner as in Experiment 2. The testing sequence was equivalent to Experiment 2, except that observers now adapted to the superimposed image of the mean gray background from the bottom monitor and the randomly chosen adapting illuminant from the top monitor.

## Acknowledgments

NIH grants EY07556 & EY13312 (QZ).

## References

1. Ferwerda JA, Pellacini F, Greenberg D (2001) A psychophysically-based model of surface gloss perception. Proc SPIE 4299:291–301.

2. Wills J, Agarwal S, Kriegman D, Belongie S (2009) Toward a perceptual space for gloss. ACM Trans Graph 28:103:1–103:15.

3. Victor JD, Conte MM (2012) Local image statistics: maximum-entropy constructions and perceptual salience. J Opt Soc Am A 29:1313–1345.

4. Lakatos S (2000) A common perceptual space for harmonic and percussive timbres. Perc & Psycho 62:1426–1439.

5. Terasawa H, Slaney M, Berger J (2005) The thirteen colors of timbre. Proc IEEE WASPAA 323–326.

6. Pols LCW, van der Kamp LJTh, Plomp R (1969) Perceptual and Physical Space of Vowel Sounds. J Acoust Soc Am 46:458–467.

7. McDermott JH, Schemitsch M, Simoncelli EP (2013) Summary statistics in auditory perception. Nat Neurosci 16:493–498.

8. Arfib D, Couturier JM, Kessous L, Verfaille V (2002) Strategies of mapping between gesture data and synthesis model parameters using perceptual spaces. Org Sound 7:127–144.

9. Giese MA, Lappe M (2002) Measurement of generalization fields for the recognition of biological motion. Vision Res 42:1847–1858.

10. Hollins M, Bensmaia S, Karlof K, Young F (2000) Individual differences in perceptual space for tactile textures: evidence from multidimensional scaling. Percep & psycho 62:1534–1544.

11. Bensmaia SJ, Denchev PV, Dammann JF, Craig JC, Hsiao SS (2008) The representation of stimulus orientation in the early stages of somatosensory processing. J Neurosci 28:776–786.

12. Cleland TA, Johnson BA, Leon M, Linster C (2007) Relational representation in the olfactory system. Procd Natl Acad Sci USA 104:1953–1958.

13. Zaidi Q, Victor J, McDermott J, Geffen M, Bensmaia S, Cleland TA (2013) Perceptual spaces: mathematical structures to neural mechanisms. J Neurosci 33:17597–17602.

14. Klein F (1939) Elementary Mathematics from an Advanced Standpoint: Geometry (Dover, New York, NY, USA).

15. Brannan DA, Esplen MF, Gray JJ (1999) Geometry (Cambridge University Press, Cambridge, UK).

16. Kirchner E (2015) Color theory and color order in medieval Islam: a review. Color Res Appl 40:5–16.

17. Smithson HE, Dinkova-Bruun G, Gasper GEM, Huxtable M, McLeish TCB, Panti C (2012) A three-dimensional color space from the 13th century. J Opt Soc Am A 29:A346–A352.

18. Maxwell JC (1860) IV. On the Theory of Cmpound Colours, and the Relations of the Colours of the Spectrum. Phil Trans R Soc Lond 150:57–84.

19. Schnapf, JL, Kraft TW, Baylor DA (1987) Spectral sensitivity of human cone photoreceptors. Nature 325:439–441.

20. Smith VC, Pokorny J (1975) Spectral sensitivity of the foveal cone photopigments between 400 nm and 500 nm. Vision Res 15:161–171.

21. MacLeod DIA, Boynton RM (1979) Chromaticity diagram showing cone excitation by stimuli of equal luminance. J Opt Soc Am 69:1183–1186.

22. D. L. MacAdam (1942) Visual Sensitivities to Color Differences in Daylight. J Opt Soc Am 32:247–273.

23. Le Grand Y (1949) Les seuils différentiels de couleurs dans la théorie de Young. Rev d’Optique Théor et Instr 28:261–278.

24. Friele LFC (1961) Analysis of the Brown and Brown-MacAdam colour discrimination data. Farbe 10:193–224.

25. Wyszecki G, Stiles WS (1982) Color science: Concepts and methods, quantitative data and formulae (John Wiley and Sons, New York, NY).

26. Krauskopf J, Gegenfurtner K (1992) Color discrimination and adaptation. Vision Res 32:2165–2175.

27. Zaidi Q (1998) Identification of illuminant and object colors: heuristic-based algorithms. J Opt Soc Am A 15(7):1767–76.

28. Zaidi Q, Bostic M (2008) Color strategies for object identification. Vision Res 48:2673–81.

29. Radonjic A, Cottaris NP, Brainard DH (2015) Color constancy supports cross-illumination color selection. J Vision 15:1–13.

30. Shepard RN (1962) The analysis of proximities: Multidimensional scaling with an unknown distance function. Part I. Psychometrika 27:125–140.

31. Indow T (1980) Gobal color metric and color-appearance systems. Color Res Appl 5:5–12.

32. Wuerger SM, Maloney LT, Krauskopf J (1995) Proximity judgments in color space: Tests of a Euclidean color geometry. Vision Res 35:827–835.

33. Coxeter HSM, Greitzer SL (1967) Quadrangles: Varignon’s Theorem in Geometry Revisited (Math Assoc Amer, Washington, DC, USA), p. 51–56.

34. Chichilnisky EJ, Wandell BA (1999) Trichromatic opponent color classification. Vision Res 39:3444–3458.

35. Smithson H, Zaidi Q (2004) Color constancy in context: Roles for local adaptation and levels of reference. J Vision 4:693–710.

36. Pinto N, Doukhan D, DiCarlo JJ, Cox DD (2009) A high-throughput screening approach to discovering good forms of biologically inspired visual representation. PLoS Comp Bio 5:e1000579.

37. Plateau JAF (1872) Sur la mesure des sensations physiques, et sur la loi qui lie l’intensité de ces sensations á l’intensité de la cause excitante. Bull Acad Roy Belg, 33:376–388.

38. Falmagne, JC (2002) Elements of psychophysical theory (Oxford University Press, Oxford, UK).

39. Goodman N (1972) Seven strictures on similarity in Problems and Projects (Bobbs-Merrill, Indianapolis, IN, USA).

40. Zaidi Q (2001) Color constancy in a rough world. Color Res Appl 26:S192–S200.

41. Wool LE, Komban SJ, Kremkow J, Jansen M, Li X, Alonso JM, Zaidi Q (2015) Salience of unique hues and implications for color theory. J Vision 15:1–11.

42. Bergström SS (1977) Common and relative components of reflected light as information about the illumination, colour, and three-dimensional form of objects. Scand J Psych 18:180–186.

43. Zaidi Q, Ennis R, Cao D, Lee B (2012) Neural locus of color afterimages. Curr Bio 22:220–224.

44. Wertheimer M (1912) Experimentelle Studien über das Sehen von Bewegung. Zeit für Psych 61:161–265.

45. Tversky A (1977) Features of similarity. Psych Rev 84:327–352.

46. Shepard, RN (1987) Toward a universal law of generalization for psychological science. Science 237:1317–1323.

47. Tennenbaum JB, Griffiths TL (2001) Generalization, similarity, and Bayesian inference. Behav Brain Sci 24:629–641.

48. Zaidi Q, Marshall J, Thoen H, Conway BR (2014) Evolution of neural computations: Mantis shrimp and human color decoding. i-Perception 5:492–496.

49. Zaidi Q, Halevy D (1993) Visual mechanisms that signal the direction of color changes. Vision Res 33:1037–1051.

50. Jain A, Zaidi Q (2013) Efficiency of extracting stereo-driven object motions. J Vision, 13:1–14.

## References

1. MacLeod DIA, Boynton RM (1979) Chromaticity diagram showing cone excitation by stimuli of equal luminance. J Opt Soc Am 69:1183–1186.

2. Derrington AM, Krauskopf J, Lennie P (1984) Chromatic mechanisms in the lateral geniculate nucleus of macaque. J Physiol 357:241–265.

3. Zaidi Q, Halevy D (1993) Visual mechanisms that signal the direction of color changes. Vision Res 33:1037–1051.

4. Jain A, Zaidi Q (2013) Efficiency of extracting stereo-driven object motions. J Vision 13:1–14.

5. Marcu G (2003) Gray tracking correction for TFT-LCDs in Proc SPIE 5293, Color Imaging IX: Processing, Hardcopy, and Applications.

6. Haenel V (2009) pyoptical: Python interface to CRS OptiCAL. Available at: https://github.com/esc/pyoptical.

